# Chromosome-level assembly of the gray fox (*Urocyon cinereoargenteus*) confirms the basal loss of *PRDM9* in Canidae

**DOI:** 10.1101/2023.11.08.566296

**Authors:** Ellie E. Armstrong, Ky L. Bissell, H. Sophia Fatima, Maya A. Heikkinen, Anika Jessup, Maryam O. Junaid, Dong H. Lee, Emily C. Lieb, Josef T. Liem, Estelle M. Martin, Mauricio Moreno, Khuslen Otgonbayar, Betsy W. Romans, Kim Royar, Mary Beth Adler, David B. Needle, Alex Harkess, Joanna L. Kelley, Jazlyn A. Mooney, Alexis M. Mychajliw

**Affiliations:** Washington State University, Pullman, WA, USA; Department of Biology, Middlebury College, Middlebury, VT, USA; Program in Environmental Studies, Middlebury College, Middlebury, VT, USA; Vermont Department of Fish and Wildlife, VT, USA; University of New Hampshire, Durham, NH, USA; HudsonAlpha Institute for Biotechnology, Huntsville, AL, USA; University of California, Santa Cruz, Santa Cruz, CA, USA; University of Southern California, Los Angeles, CA, USA

**Keywords:** Gray fox, canidae, PRDM9, heterozygosity

## Abstract

Reference genome assemblies have been created from multiple lineages within the Canidae family, however, despite its phylogenetic relevance as a basal genus within the clade, there is currently no reference genome for the gray fox (*Urocyon cinereoargenteus*). Here, we present a chromosome-level assembly for the gray fox (*U. cinereoargenteus*), which represents the most contiguous, non-domestic canid reference genome available to date, with 90% of the genome contained in just 34 scaffolds and a contig N50 and scaffold N50 of 59.4 and 72.9 Megabases (Mb), respectively. Repeat analyses identified an increased number of simple repeats relative to other canids. Based on mitochondrial DNA, our Vermont sample clusters with other gray fox samples from the northeastern US and contains slightly lower levels of heterozygosity than gray foxes on the west coast of California. This new assembly lays the groundwork for future studies to describe past and present population dynamics, including the delineation of evolutionarily significant units of management relevance. Importantly, the phylogenetic position of *Urocyon* allows us to verify the loss of *PRDM9* functionality in the basal canid lineage, confirming that pseudogenization occurred at least 10 million years ago.

## Introduction

Reference genomes have become valuable tools in conservation science and decision-making (Supple and Shapiro 2018; Formenti et al. 2022; Paez et al. 2022). While mammals tend to be the most well-represented amongst large-scale consortia endeavors (e.g., Zoonomia, Zoonomia Consortium 2020), progress on generating data for the mammalian family Canidae has lagged, with only six of the thirteen extant genera having publicly available, representative reference genome assemblies. Canidae currently contains 39 extant species that vary in size, ecology, and distribution, and diverged from other carnivoran families approximately 40-60 million years ago (Wayne 1993; Nyakatura and Bininda-Emonds 2012).

Within Canidae, the genus *Urocyon* has historically been difficult to place, but is thought to represent the sister lineage to all other living canids (Tedford et al. 1995; Lindblad-Toh et al. 2005; Nyakatura and Bininda-Emonds 2012). The genus contains only two species: the gray fox (*Urocyon cinereoargenteus,* Schreber 1775), which is found from southern Canada through northern South America, and the island fox (*Urocyon littoralis*), which is restricted to the California Channel Islands. Gray foxes are grizzled in appearance and have a number of scansorial (climbing) adaptations that facilitate their use of deciduous forests (Fritzell and Haroldson 1982). However, there is significant variation in their responses to habitat alterations across their North American range, with some studies suggesting they are tolerant of habitat disturbance and extremely abundant, while others suggest they are highly sensitive and occur at lower population densities (McAlpine et al. 2008; Bauder et al. 2020; Allen et al. 2021).

The lack of precise ecological knowledge concerning the gray fox across its range is compounded by uncertainty in the species’ historical distribution and genetic structure, and the perceived expansion of gray foxes into urban ecosystems and new geographic areas is accompanied by the potential for human-wildlife conflict and disease spillover. For example, a gray fox in New England was diagnosed with concurrent infections of antibiotic resistant *Listeria monocytogenes*, skunk adenovirus-1, and canine distemper virus (Needle et al. 2020). Both mitogenomes and genotyping by sequencing approaches suggest that western and eastern North American gray fox populations diverged in the early-mid Pleistocene (∼0.8 million years ago) and now form a secondary contact zone of eastern and western lineages at the Great Plains Suture (Reding et al. 2021; Kierepka et al. 2023), displaying a distinct pattern from many other North American carnivores, and disagreeing with previously held morphological subspecies designations. Eastern populations of the gray fox are phylogenetically distinct from all other gray fox populations, and the island fox (*U. littoralis*) is nested within the western gray fox clade (Preckler-Quisquater et al. 2023). This cryptic divergence pattern, coupled with recent heterogenous range expansions and population declines (e.g., McAlpine et al. 2008), underscores the need for additional genomic resources for this species.

The gene PRDM9 directs the majority of meiotic recombination events in humans and most mammals by directing the location of double stranded breaks in the genome (Baudat et al. 2010; Myers et al. 2010; Parvanov et al. 2010). Nevertheless, PRDM9 is pseudogenized across wild (e.g. Ethiopian wolf, red fox, coyote, and dhole) and domestic canids, (Munoz-Fuentes et al. 2011; Axelsson et al. 2012; Mooney et al. 2013) as well as other vertebrate species. Though the mechanisms that direct recombination in canids are unknown, most recombination events tend to be directed towards promoters and GC-rich regions (Auton et al. 2013). More recent work has posited that the loss occurred in the branch leading to Canidae, approximately 14-46 million years ago (Cavassim et al. 2022). However, the authors did not have access to sequence data from the most basal canid lineage (*U. cinereoargenteus*) to verify the loss. Given the high-quality of our new reference assembly, we sought determine whether the pseudogenization of PRDM9 occurred before or after the differentiation of Canidae. We built a chromosome-level reference assembly for gray fox from a male individual from Vermont, which can be used as a foundation to answer pressing questions about the phylogenetic relevance of *Urocyon* as a basal genus within Canidae, its importance in a One Health context, the uncertainty regarding the antiquity of the island fox (Hofman et al. 2016; Sacks et al. 2022), and whether the Eastern and Western clades of North American gray fox may represent distinct species (Hofman et al. 2015; Goddard et al. 2015; Reding et al. 2021). We assessed the quality of our assembly relative to other available canid genomes, contextualized our mitogenome and heterozygosity within available data for *Urocyon*, and assessed the functionality of the PRDM9 gene. This reference genome expands the possibilities for future studies of gray fox chromosomal architecture, population-level diversity, and disease ecology.

## Methods & Materials

### Sample collection, sequencing, and assembly

In October 2021, the Vermont Department of Fish and Wildlife obtained a liver sample from a deceased adult male gray fox (*U. cinereoargenteus*) donated by licensed fur trappers as part of ongoing wildlife health surveillance studies (Figure 1). The sample was shipped on wet ice to Cantata Bio (Santa Cruz, CA, USA). Liver tissue (100mg) was combined with 10 mls of G2 buffer + Rnase producing 60ug of DNA after incubation. A maxi column was used to spin the DNA down, followed by a wash with ethanol. The resulting pellet of DNA was dissolved in TE.

**Figure 1:**
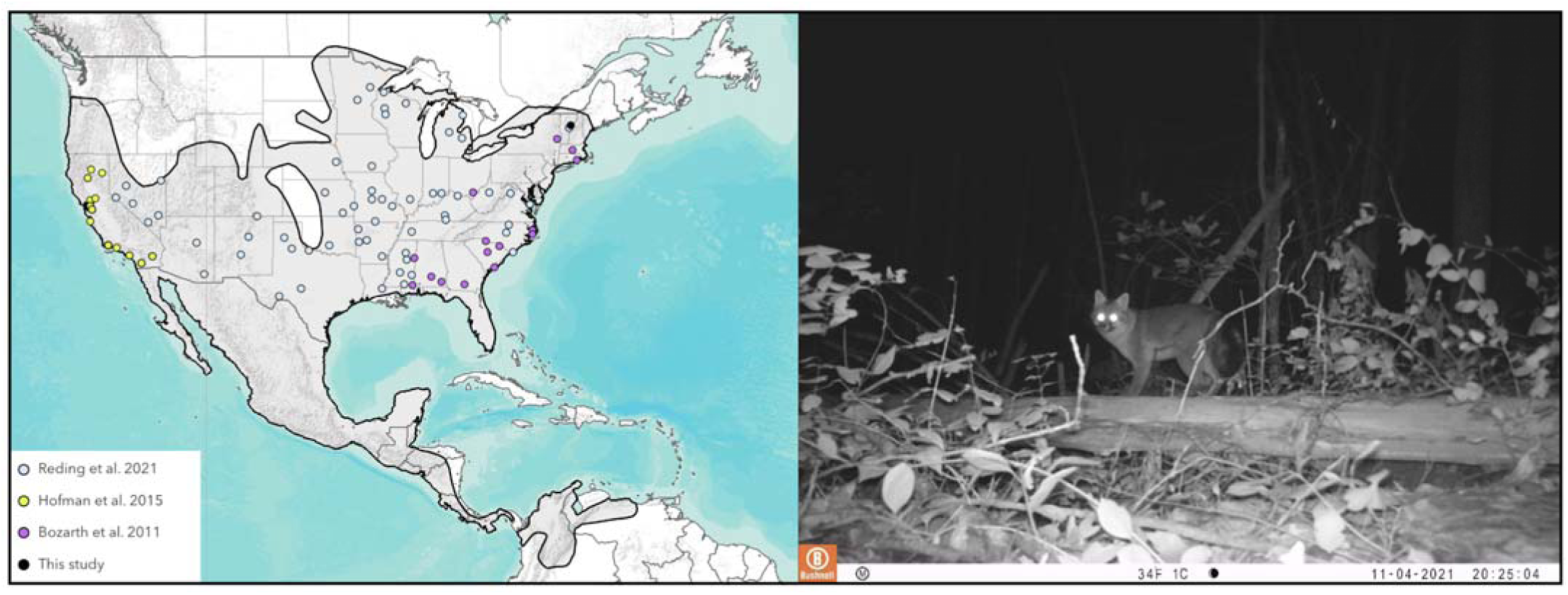
Left: Map showing the location of gray fox samples used in some genetic studies and the location of our new reference genome from Vermont, overlaid on the current gray fox range (IUCN). Map sources: Esri, USGS, NOAA. Right: A gray fox explores a commonly used hiking trail on the Middlebury College campus (photo by Andrew Ng, Middlebury College).

DNA samples were quantified using Qubit 2.0 Fluorometer (Life Technologies, Carlsbad, CA, USA). The PacBio SMRTbell library (∼20kb) for PacBio Sequel was constructed using the SMRTbell Express Template Prep Kit 2.0 (Pacific Biosciences, Menlo Park, CA, USA) following the manufacturer recommended protocol. The library was bound to polymerase using the Sequel II Binding Kit 2.0 (PacBio) and loaded onto the PacBio Sequel II. Sequencing was performed on PacBio Sequel II 8M SMRT cells.

PacBio CCS (124 Gb in total) reads were used as an input to Hifiasm v0.15.4-r347 (Cheng et al. 2021) with default parameters. BLAST (ncbi-blast+/2.11.0; Camacho et al. 2009) results of the Hifiasm output assembly (hifiasm.p_ctg.fa) against the nucleotide database were used as input for blobtools2 v1.1.1 (Laetsch and Blaxter 2017) and contigs identified as possible contamination were removed from the assembly (filtered.asm.cns.fa). Finally, purge_dups v1.2.5 (Guan et al. 2020) was used to remove haplotigs and contig overlaps (purged.fa).

Chromatin from liver samples was fixed in place with formaldehyde in the nucleus and then extracted to construct each Dovetail Omni-C library. Fixed chromatin was digested with DNAse I, chromatin ends were repaired and ligated to a biotinylated bridge adapter followed by proximity ligation of adapter containing ends. After proximity ligation, crosslinks were reversed and the DNA was purified. Purified DNA was treated to remove biotin that was not internal to ligated fragments. Sequencing libraries were generated using NEBNext Ultra enzymes and Illumina-compatible adapters. Biotin-containing fragments were isolated using streptavidin beads before PCR enrichment of each library. The library was sequenced on an Illumina HiSeqX platform to produce ∼ 30x sequence coverage.

The *de novo* assembly produced by Hifiasm and Dovetail OmniC library reads were used as input data for HiRise (accessed September 2022), a software pipeline designed specifically for using proximity ligation data to scaffold genome assemblies (Putnam et al. 2016). Dovetail OmniC library sequences were aligned to the draft input assembly using BWA-MEM v0.7.17-r1188 (Li and Durbin 2009) The separations of Dovetail OmniC read pairs mapped (MQ >50) within draft scaffolds were analyzed by HiRise to produce a likelihood model for genomic distance between read pairs, and the model was used to identify and break putative misjoins, to score prospective joins, and make joins above a threshold.

### Assembly quality and continuity

To assess the size distribution of contigs and scaffolds, as well as the quality and continuity of our genome assembly, we used scripts from Assemblathon2 (Bradnam et al. 2013). We compared our assembly to assemblies of other representative canids with available genome assemblies (Table 1).

**Table 1.**
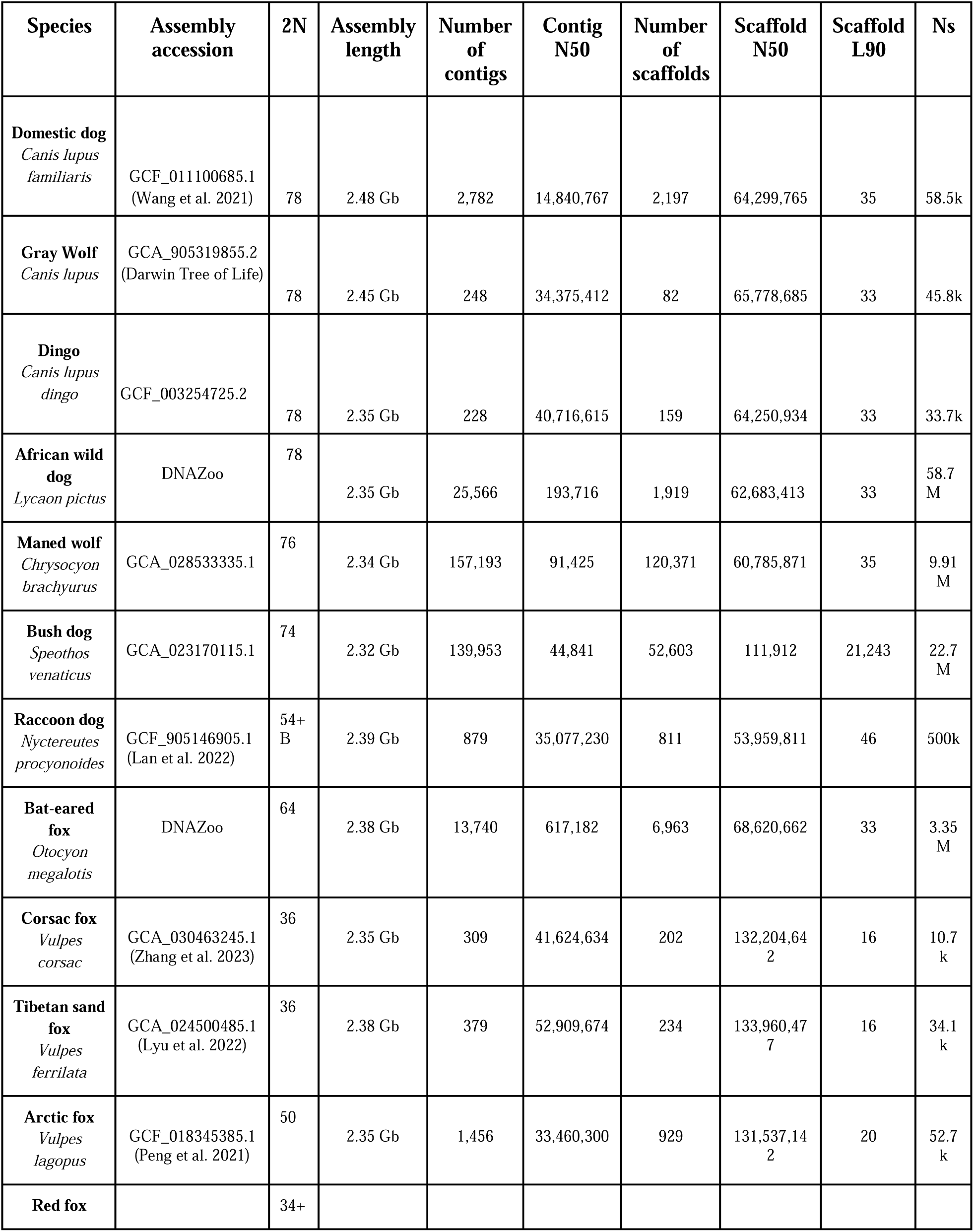
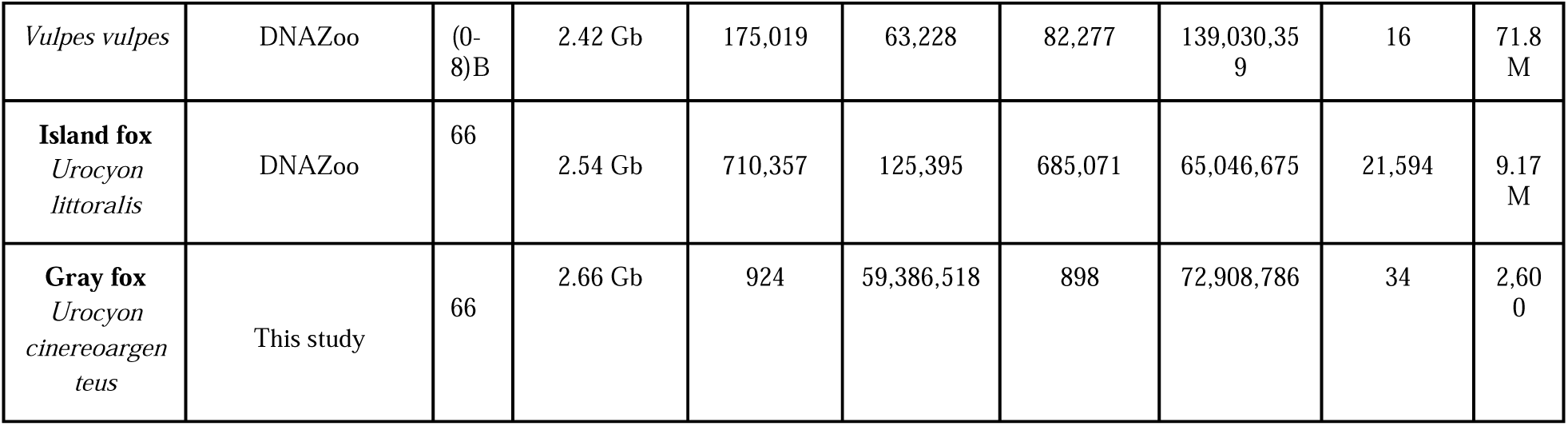
Assembly statistics for representative published Canidae assemblies and the gray fox genome from this study.

We assessed the completeness of genes using the compleasm v0.2.2 program (Huang and Li 2023), which is a faster application of BUSCO (Simão et al. 2015; Waterhouse et al. 2018). Compleasm leverages the BUSCO framework to search genomes for highly conserved sets of orthologs and evaluates whether they are present in the genome in a single copy, or are duplicated, fragmented, or missing. We utilized the provided carnivora_odb10 gene set to query the canid genomes.

Finally, we investigated chromosomal rearrangements and chromosomal contiguity. We aligned the genomic sequence of the Gray fox to the Arctic fox (GCA_018345385; Peng et al. 2021) and the domestic dog (GCF_011100685; Wang et al. 2021) using minimap2 v2.24 (Li 2018), followed by visualization with CIRCA v1.2.3 software (https://omgenomics.com/circa/), which allows for easy visualization of syntenic regions between the genomes. Prior to plotting, we used the pafr program v0.0.2 (https://github.com/dwinter/pafr) to read in and filter PAF files and subsequently convert them to CSV in R v4.2.1 (R Core Team 2021). We discarded alignments with a length less than 25,000 bp and a map quality less than 60. Alignments were also restricted to autosomes and the X chromosome, as some assemblies were female and others did not have identified Y chromosomes.

### Mitochondrial phylogenetics

We used minimap2 to align the PacBio raw reads from *U. cinereoargenteus* against a reference mitogenome (MW600067.1; Reding et al. 2021).The resulting SAM file was then converted to a BAM file using SAMtools v1.16.1 (Li et al. 2009; Danecek et al. 2021), and a consensus sequence called using ANGSD v0.940 (Korneliussen et al. 2014) with the flags “-doFasta 2” and “-doCounts 1” to produce a consensus mitochondrial genome for the fox. Then, existing complete mitochondrial genome data from *U. cinereoargenteus* was downloaded from NCBI genbank (N = 102, NCBI Popset 2239967418 and 765367839; Hofman et al. 2015; Reding et al. 2021). The FASTA files were concatenated alongside the consensus mitogenome of the gray fox sampled in this study. The resulting FASTA file underwent multiple sequence alignment using the MAFFT v7.515 program (Katoh and Standley 2013). The output was an alignment file (.aln) which was used to generate a Bayesian phylogenetic tree with the outgroup as the root of the tree using IQ-TREE v1.6.12 (Minh et al. 2020) with flags ‘-b 100’ and ‘-m TEST’, which run 100 bootstrap replicates and find the best nucleotide substitution model, respectively. The tree was subsequently visualized using FigTree v1.4.4 (Rambaut 2007), rooted on the Western clade, and colored according to region of origin.

### Heterozygosity

All additional *U. cinereoargenteus* and *U. littoralis* whole-genome sequences available at the time of this study (January 2023) were obtained from NCBI (Robinson et al. 2016; Robinson et al. 2018) bioprojects PRJNA312115 and PRJNA478450. Short-read data was mapped to the gray fox genome assembly using BWA-MEM and converted to BAM format, sorted, indexed using SAMtools. We estimated the site frequency spectrum (SFS) using ANGSD by inputting the BAM files, along with flags ‘-anc’, ‘-ref’, ‘-fold 1’, ‘-dosaf 1’, ‘-GL 2’, ‘-C 50’, ‘-minq 20’, and ‘-minmapq 30’. The reference gray fox genome was provided for the ‘-anc’ and ‘-ref’ flags. The ‘-fold’ flag was used to assign the reference sequence as the ancestral allele and fold the site frequency spectrum. Flags ‘-C 50’, ‘-minq 20’, and ‘-minmapq 30’ were used to remove reads and bases with low mapping and genotype call qualities. Subsequently, we ran realSFS (within ANGSD) and estimated the number of heterozygotes using the folded spectra in compliance with ANGSD guidelines (see http://www.popgen.dk/angsd/index.php/Heterozygosity).

### Repetitive elements

We estimated repeat content in our *de novo* gray fox assembly as well as other representative canid genomes (Table 1) using TETools v1.7 (Lerat et al. 2017). For each genome, we first built a database using the *BuildDatabase* command. Subsequently, we used the *RepeatModeler* command, which uses the RepeatModeler v2.0.2 tool (Flynn et al. 2020) to locate and annotate repeats *de novo* in sequences and genome assemblies. Last, we used the command *RepeatMasker* v4.1.4 (Flynn et al. 2020) within TETools, which takes the output from RepeatModeler, performs additional screening for repeats within the genome based on homology using the dfam database v3.6 (Storer et al. 2021), and outputs a summary of the repeats in the input sequence.

### Functionality of PRDM9

We used Seqtk v1.3-r117 (https://github.com/lh3/seqtk) to extract the human PRDM9 gene sequence from the composite sequence dataset used in (Mooney et al. 2023). Then, we used this human PRDM9 sequence data to locate the PRDM9 ortholog within the gray fox genome using BLAST (ncbi-blast+ v2.7.1). We identified the likely start and end of the PRDM9 gene within the gray fox genome and used Seqtk to extract this sequence, adding approximately 10k base pairs of buffer on either side to ensure that the totality of the gene sequence was extracted. Then, we concatenated the gray fox PRDM9-like sequence back into a dataset from (Mooney et al. 2023), which contained PRDM9 sequences from the human, Ethiopian wolf (*Canis simensis*), two populations of gray wolf (Arctic wolf, Isle Royale wolf; *Canis lupus*), and a number of domestic dog breeds including pug, labrador retriever, tibetan mastiff, and border collie. We used MAFFT to align all sequences from the new concatenated file, searching for any loss-of-function mutations in our alignment to reveal information about the functionality of *PRDM9* in gray foxes. Lastly, GeneWise <u>(Birney et al. 2004,</u> https://www.ebi.ac.uk/Tools/psa/genewise/) was used to align the DNA sequence from gray fox to human *PRDM9* protein sequence, which allowed us to visually identify any insertions or frameshift errors (see Supplementary Figure 1). We uploaded a merged file with all nucleotide sequences that were used to Github.

## Results & Discussion

### Assembly quality and continuity

We assembled a genome for a male gray fox using PacBio HiFi and Dovetail OmniC data. The final assembly totaled 2,658,766,243 base pairs (bp) in length across 898 scaffolds. The contig and scaffold L90 were 51 and 34, and the contig and scaffold N50 were 59.4 Mb and 72.9 Mb, respectively (Table 1). Given the statistics of the final assembly, the gray fox genome was of notably better or of equal quality compared to the other canid assemblies (Table 1), despite karyotypic differences. The scaffold L90 was close to the total number of chromosomes expected in the gray fox (32 autosomes and an X and Y chromosome; Graphodatsky et al. 2008), indicating that it is likely that most autosomes and the X chromosome are contained in approximately one scaffold.

We identified linkage groups using whole-genome alignment in conjunction with information obtained from previous physical mapping studies (Graphodatsky et al. 2008). Whole-genome alignments confirmed that the expected linkage groups for all autosomes and the X chromosome were present (Figure 2). As the Y chromosome is difficult to assemble due to its repetitive nature (Burgoyne 1982), we did not identify any Y chromosome scaffolds or contigs as part of this work; there are only limited and incomplete representations of canid Y chromosomes that are publicly available. However, we were able to observe multiple autosomal fissions and fusions relative to the domestic dog karyotype which have been previously identified in karyotypic studies (Figure 2; Nie et al. 2012; Perelman et al. 2012).

**Figure 2.**
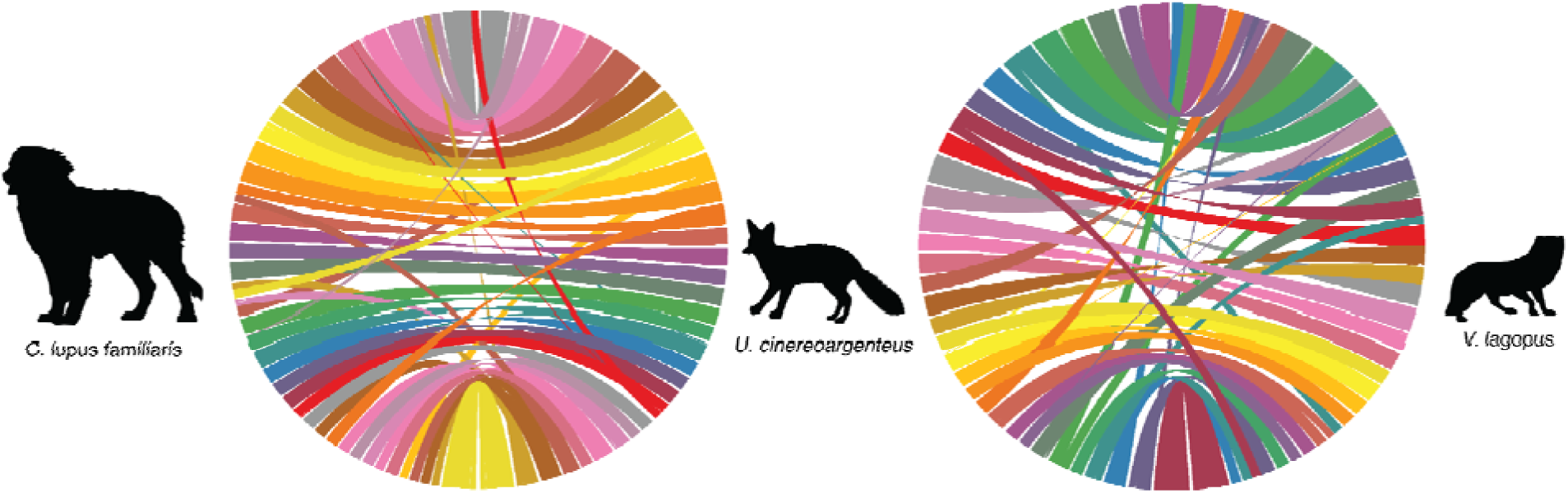
Whole-genome alignments for (A) the domestic dog (GCF_011100685.1) and gray fox and (B) the arctic fox (GCF_018345385.1) and the gray fox. Colored ribbons originate on the gray fox chromosomes (center), so that each unique color is assigned to one gray fox chromosome. In each alignment, the X chromosome is positioned at the bottom of the alignment.

We also assessed the quality of the genome assembly using compleasm (Huang and Li 2023). Broadly, canid assemblies scored relatively high with most assemblies having over 90% of the genes queried in complete and single copies, with the exception of the African wild dog and the bush dog (Table 2). The gray fox genome assembled here had the highest single-copy score of any canid genome assembly assessed, which confirms that most gene regions were assembled well.

**Table 2:**
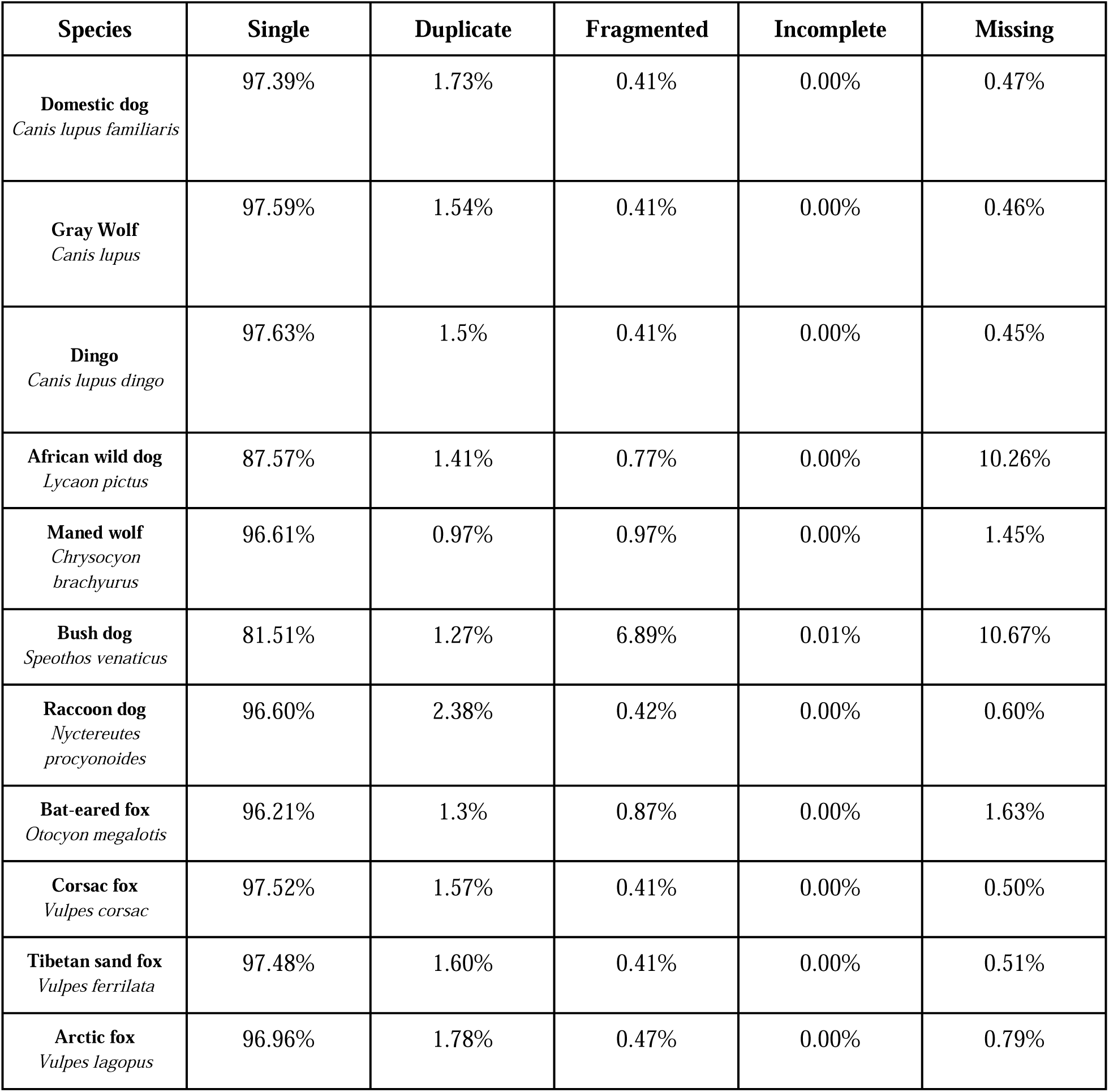
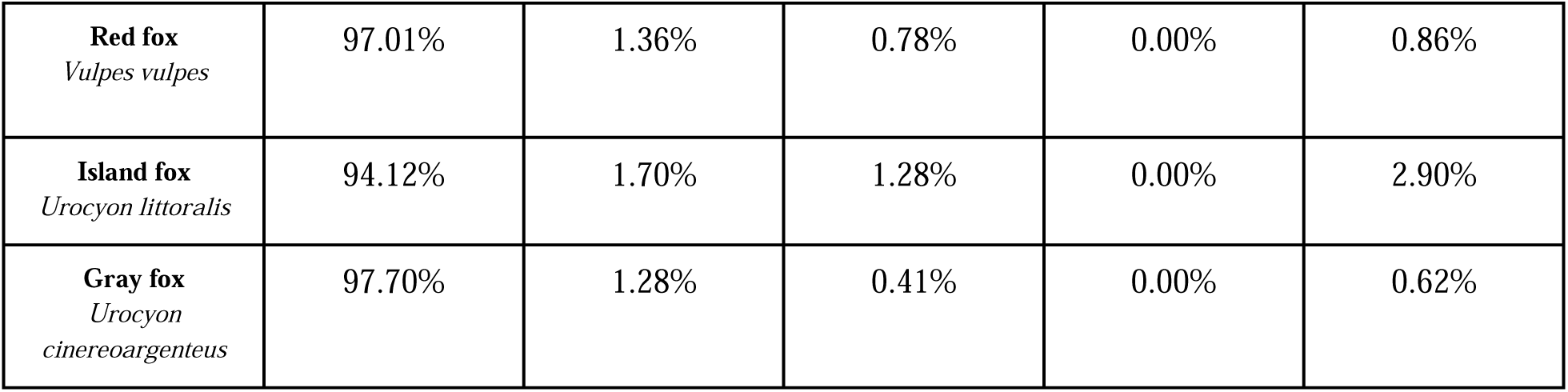
Compleasm results for representative published Canidae assemblies and the gray fox genome from this study.

### Repeat Elements

We found that the quantity and classification of repeat elements were relatively similar across the Canidae family. However, gray foxes – but not island foxes – have an increased number of simple sequence repeats (SSRs) compared to Arctic foxes, dogs, gray wolves, and red foxes (Figure 3). SSRs, also known as microsatellites, are short repeating motifs of six base pairs or less. SSRs are generally thought to evolve through the process of slippage (Schlötterer and Tautz 1992; Ellegren 2004) and have been broadly leveraged to query the diversity and identity of individuals across the tree of life through microsatellite typing (Wright and Bentzen 1995; Schlötterer and Pemberton 1994). They have been implicated in many processes, such as transcription regulation (Kashi and King 2006; Hancock and Simon 2005) and disease (Hancock and Simon 2005), and expansions of these elements may also contribute broadly to genomic instability (Khristich and Mirkin 2020).

**Figure 3:**
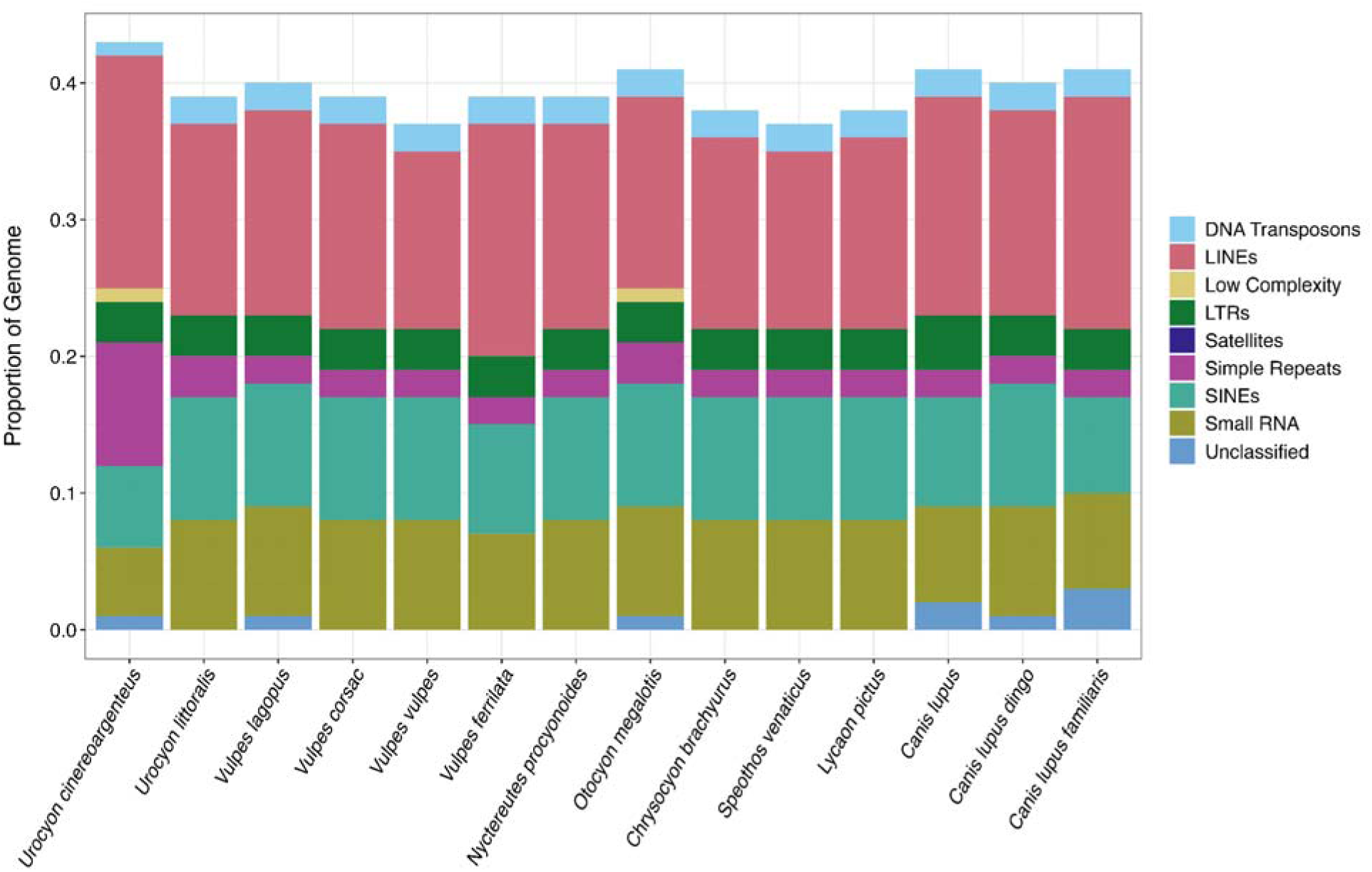
Repeat element proportions for members of the *Canidae* family with reference genome assemblies. Species are ordered phylogenetically. Proportions correspond to the proportion of base pairs in each repeat category as identified by TETools.

It is unclear whether the increased number of SSRs in the gray fox is the ancestral state and SSRs were subsequently lost in the remainder of the canids, or whether the expansion was more recent, at least after the establishment of the western gray fox or island fox lineage. Additionally, the island fox genome assembly, despite being in chromosomes, has a very low contig N50 since it was assembled using short read data (Table 1). It is thus unclear if the absence of SSRs in the island fox genome assembly (or the difference relative to what we found in the gray fox) is due to an assembly artifact. However, previous studies comparing genomes which have been assembled using short-read data (similar to the island fox genome assembly) have in general suggested that even highly fragmented assemblies will yield accurate repeat content statistics (Armstrong et al. 2020). Additional investigation into the timing of the SSR expansion, and whether it is indeed absent in the island species, may help to elucidate the timing of diversification of the lineages and provide information on the canid ancestral genome.

### Fossil record, range, and phylogeographic patterns

The genus *Urocyon* extends back into the Pliocene (Hemiphillian North American Land Mammal Age) (Kurten & Anderson 1980). Shifts in glacial/interglacial cycles (e.g., Sangamonian, Wisconsin) could have resulted in range expansions and contractions, events which could be used to test contemporary genomic structure and dispersal patterns. For example, our observations of heterozygosity (see below) are consistent with the hypothesis that there was a recent expansion of gray foxes from a southern refugium into the northeast post-Pleistocene (Bozarth et al 2011), however additional samples from populations in the eastern and northeastern U.S. have not been explored, even in recent papers which generated genome-level population data for the species (Kierepka et al. 2023; Preckler-Quisquater et al. 2023).

Our mitochondrial phylogeny recovered a division between the eastern and western haplotypes of gray fox, consistent with previous results (Goddard et al. 2015; Reding et al. 2021; Kierepka et al. 2023; Figure 4). The consensus mitochondrial genome for the Vermont gray fox sample fell within the eastern clade and clustered with the single other Vermont mitochondrial sample available from the literature (Figure 4). We observed no intermixing between samples originating from the eastern and western United States, supporting previous evidence that admixture between groups is likely limited (Bozarth et al. 2011; Reding et al. 2021). In several recent studies, novel genomic data revealed a contact zone between the eastern and western clades in the plains region in Texas and Oklahoma (Kierepka et al. 2023, Preckler-Quisquater et al. 2023). These studies did not contain samples from most of the northeastern range of the foxes outside of some samples in Tennessee and South Carolina, which will be important in understanding the range wide diversity and structure of the gray fox. Future analyses using this reference genome, combined with a reference genome recently released for the Santa Catalina island fox (*U. littoralis catalinae*) (Hendricks et al. 2022), will be important for understanding the taxonomic relationship between these two lineages, with consequences for species delimitation and management.

**Figure 4.**
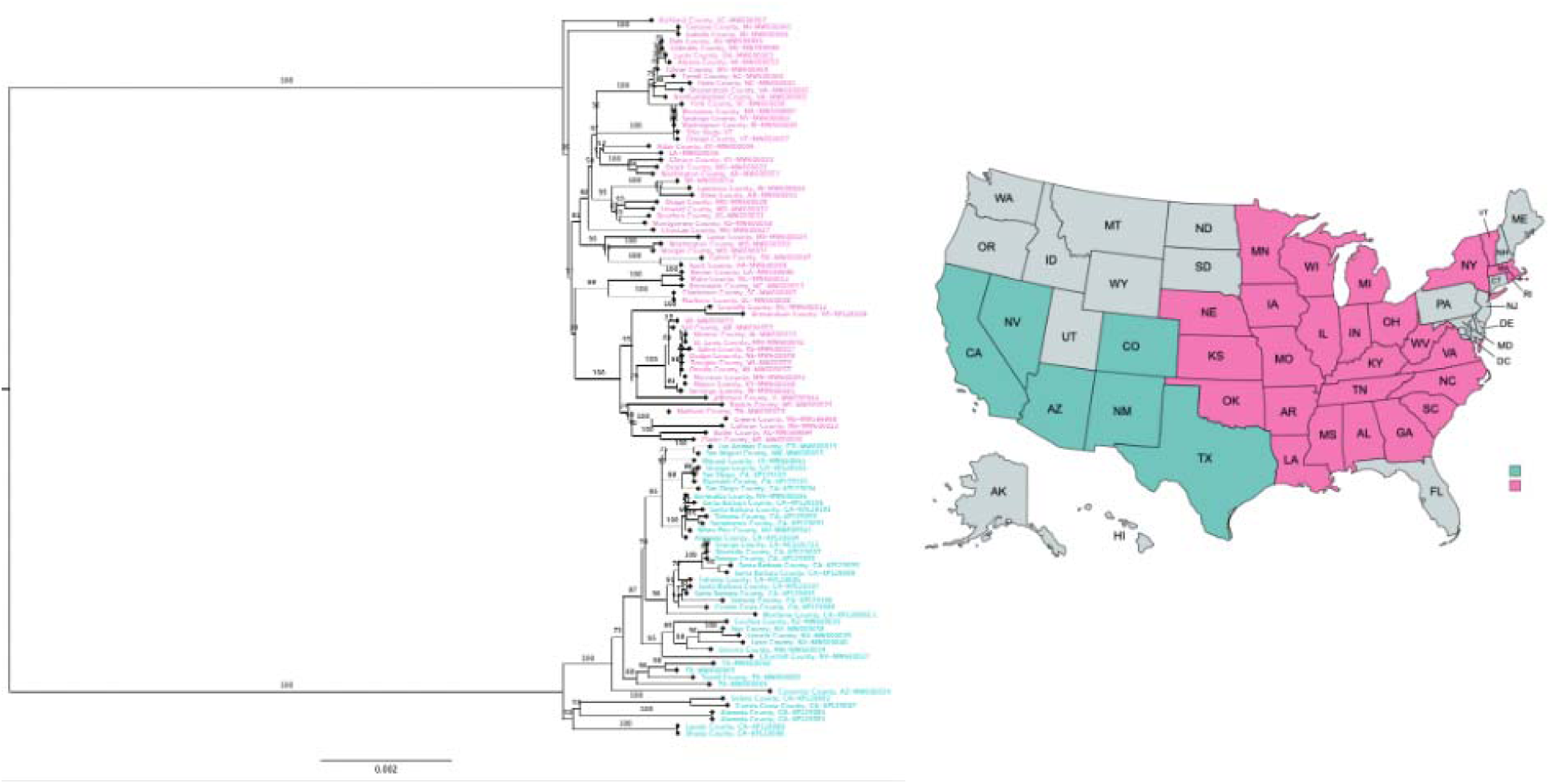
Phylogenetic tree generated from an analysis of 104 unique Urocyon *cinereoargenteus* mitogenome haplotypes. Coloring of the sample codes reflect the location origin of the sample. Turquoise=Western United States (N=42), Pink=Eastern United States (N=61).

The natural history literature contains a number of contrasting statements regarding the baseline range of gray foxes in the northeastern United States, particularly New England. Some have hypothesized that gray fox populations declined extensively in the 19^th^ century, and has since been followed by expansions in the past 50 years linked to a warming climate (Finley & Godin 1978; Banfield 1974). Recent genomic analyses did not investigate the most northeastern parts of the gray fox range, thus overlooking archaeological and paleontological records from this region… For example, in 1635 CE, pilgrims in Massachusetts described “two or three kinds of fox, one a great yellow Fox, another Grey, who will climb up into trees” (Keay 1901) and is known from the pre-contact zooarchaeological record of Martha’s Vineyard (∼400-1100 years before present) (Huntington 1959). By 1931 CE, the “Mammals of Hampshire County, Massachusetts” noted that the species is common, but was perceived as less common than the red fox because of its dense forest association and less desirable fur (Crane 1931).

However, not all areas of New England have a long record of gray fox presence. Moving northward, the “Notes on New Hampshire Mammals” does not include the gray fox in the list of native mammals occurring from 1915-1920 CE (Jackson 1922). The earliest gray fox pelt in a museum collection from Vermont is from 1910 CE (MCZ:Mamm:64310). Osgood (1938) noted in “The Mammals of Vermont” that the subspecies *U. c. borealis* “reaches its northern limit in Vermont”, with the farthest north occurrence considered in Whiting (Champlain Valley) in western Vermont and Woodstock in eastern Vermont, and by 1938, reported confirmed skulls from Rutland and Springfield. In contrast with Massachusetts, the gray fox is not present in the zooarchaeological record of Vermont, despite the Holocene presence of other carnivores such as red foxes, fishers, and martens (Mychajliw et al. 2023). The hypothesis that gray foxes are a relatively new addition to the canid community of Vermont, starting in the early 20^th^ century, has yet to be tested with genomic data.

### Heterozygosity

The Vermont gray fox had slightly lower levels of heterozygosity compared to those known from California (Table 3) Congruent with previous results (Robinson et al. 2016; Robinson et al. 2018), we found that gray foxes exhibited higher levels of heterozygosity than island foxes restricted to the California Channel Islands (*U. littoralis*) (Table 3). These results are consistent with a recent study showing that gray fox populations east of the contact zone had significantly lower heterozygosity than foxes from areas to the west of the contact zone (Preckler-Quisquater et al. 2023) and that diversity decreased with increasing latitude (Kierepka et al. 2023). Our preliminary results here suggest that diversity may only be slightly lower at these expansion fronts. Though our sample sizes limit our ability to make inferences about the patterns observed, they reinforce the need for sampling gray foxes across their northern and eastern range extents to fully reconstruct possible ancient refugia and recent expansion patterns.

**Table 3:** Mean heterozgyosity of gray (*U. cinereoargenteus*) and island fox (*U. littoralis*) as estimated in ANGSD.

| **Species** | **Location** | **Sample size** | **Mean heterozygosity per base pair** | **Interquartile range** |
| --- | --- | --- | --- | --- |
| *U. cinereoargenteus* | Vermont | 1 | 0.002126 | N/A |
| *U. cinereoargenteus* | California | 2 | 0.002277 | 8.315e-07 |
| *U. littoralis* | San Miguel Island | 2 | 0.000417 | 5.55805e-05 |
| *U. littoralis* | Santa Rosa Island | 3 | 0.001840 | 0.001684175 |
| *U. littoralis* | Santa Cruz Island | 2 | 0.000835 | 3.108e-06 |
| *U. littoralis* | San Nicolas Island | 3 | 0.000635 | 0.000342932 |
| *U. littoralis* | Santa Catalina Island | 2 | 0.001333 | 0.000163839 |
| *U. littoralis* | San Clemente Island | 2 | 0.000655 | 0.0001764175 |

### Functionality of PRDM9

Using our new reference genome, we identified four frameshift mutations and an additional stop codon in the zinc-finger region of *PRDM9* in the gray fox using GeneWise (Supplementary Figure 1). Recent work on examining *PRMD9* functionality demonstrated that *PRDM9* function was lost across all wolf-like canids, including both the Ethiopian wolf and Dhole (Mooney et al. 2023), but the loss had yet to be confirmed in other canids, including the gray fox. Prior to this work, Axelson and colleagues had postulated that PRDM9 was psuedogenized across all of Canidae, but their study lacked sequence data from the most divergent lineage, *Urocyon* (Axelsson et al. 2012). Our data contribute to the existing strong evidence that PRDM9 pseudogenization occurred at least 9-10MYA prior to the divergence of *Urocyon* and the remainder of the modern canid family (Eizirik et al. 2010; Lindblad-Toh et al. 2005; Matzke and Wright 2016).

### Relevance to Conservation

In the United States, the gray fox (*U. cineoargenteus*) is managed as a furbearing mammal and is regularly harvested in many states. Within Vermont, gray foxes are listed as an S5 common species with both open hunting and trapping seasons in the fall-winter, though they are not as commonly trapped as other canids. The gray fox is also involved in human-wildlife conflict: over 1,000 gray foxes across the U.S. were killed/euthanized by the USDA APHIS wildlife services program in 2022 alone (Program Data Report G, 2022). Given their expansion into suburban and urban areas and co-occurrence with multiple canids including domestic dogs, gray foxes may serve as models to understand the ecology and genomic basis of virus transmission and disease susceptibility in mesocarnivores (Henn et al. 2007). For example, a distinct clade of canine distemper virus in New England is now shared across multiple carnivore species, including the gray fox (Needle et al. 2019).

The gray fox’s wide geographic range and apparent population stability is contrasted by reports of regional declines in the Midwest (Bluett 2006; Willingham 2008; Alessi et al. 2012) and inconsistencies in fur harvest records (Bauder et al. 2020). A recent petition called for the listing of the prairie gray fox subspecies (though, we note prior that these subspecies designations are not supported by genetic data), *U. c. ocythous*, under the Endangered Species Act because of declines across its range in Iowa, Arkansas, Missouri, and Minnesota (https://www.fws.gov/species-publication-action/90-day-finding-petition-list-prairie-gray-fox-plains-spotted-skunk-and) (Department of the Interior, 2012). A dearth of data across their range, both at comparable scales and types, hinders our ability to contextualize regional patterns to see a larger picture for this species (Allen et al. 2021). Such regional declines may be driven by interference competition and intraguild predation with expanding coyotes (Egan et al. 2021). Gray foxes may have different tolerances to human activities and landscape alterations, with the advantage of a generalist diet outweighed by its more specific habitat needs (Morin et al. 2022), and may be extirpated where it cannot shift its spatial resource use (Levi and Wilmers 2012). The presence of domestic dogs may also exacerbate this competitive tension, as gray foxes shift their diel activity patterns and decrease in abundance where dogs are present (Royle and Nichols 2003). Tree cover is likely important for facilitating coexistence with increasing coyote populations, particularly in suburban areas. Parsons et al. (2022) suggest a management benchmark of 50% forest cover in a 1km radius to allow for coexistence of gray foxes and coyotes (Parsons et al. 2022).

The gray fox is currently listed by the International Union for the Conservation of Nature (IUCN) as Least Concern (Roemer et al. 2016), but such a listing could change if taxonomic revisions are made based on additional genomic research. Already, the 0.8 million year divergence timing of eastern and western gray fox clades is deeper than most intraspecific splits of other North American carnivores, such as black bears (Puckett et al. 2015). This divergence time exceeds that of interspecific splits within multiple genera in Carnivora, including the gray fox and Channel Island fox (Hofman et al. 2016; Sacks et al. 2022), as well as within two genera of South American canids, the genus *Dusicyon* (including the Falkland Islands wolf) (Austin et al 2013), and the genus *Lycalopex* (including the culpeo and Darwin’s fox) (Favarini et al. 2022). The gray fox has 16 subspecies described based on morphological features (Fritzell and Haroldson 1982), but mitochondrial haplotype data has consistently disagreed with these subspecies designations, instead suggesting more cryptic divergence patterns (Reding et al. 2021) and the potential for management unit revaluation under legislation such as the Endangered Species Act in the United States. Within Canada (where gray foxes are thought to have recently expanded), the species is currently listed as threatened under the federal Species at Risk Act,In fact, Pelee Island, Ontario, contains Canada’s only breeding population, with fewer than 300 individuals (McAlpine et al. 2008). Our reference genome provides an important tool in facilitating whole-genome, nuclear studies to resolve discrepancies between morphological and mitochondrial DNA to the benefit of conserving and managing this species across its range.

## Acknowledgements

We would like to thank David Guertin for assistance with the high-performance computing cluster (‘Ada’) at Middlebury College; this material is based upon work supported by the National Science Foundation under Grant No. 1827373. Support for this project was provided by Revive and Restore, Cantata Bio, and Dovetail Genomics as part of an ‘AG4’ program award to EEA, JLK, and AMM. We thank M. Daly and T. Swale of Cantata Bio for their contributions and facilitation of this work. We thank Dave Allen and the Middlebury Department of Biology for facilitating the Praxis course for January term students. Last, we are indebted to Alex Feltus, David Clark, and the Praxis AI team for support during the course. Figure 2 depiction of gray fox created by Gabriela Palomo-Munoz (CC BY-NC 3.0), courtesy of phylopic.org.

## Data Accessibility

All code used to process the genome can be found at https://github.com/ellieearmstrong/GrayFox_Middlebury/. Raw reads and assembly files are available under NCBI Bioproject PRJNA1005958. *The authors affirm that all data necessary for confirming the conclusions of the article are present within the article, figures, and tables*.

## Author Contributions

EMM, JTL, ECL, HSF, KLB, MM, AJ, WHL, BWR, MAH, KO, and MAJ performed research and analyzed data with the guidance of EEA and AH. JAM provided data and advice for PRDM9 analyses. EEA, JAM, and AMM designed research, performed research, analyzed data, and wrote the paper. KR, MBA, and DN contributed samples and provided guidance on the project and JLK provided guidance on the project.

