## Supplementary File 1 for "Chromosome-level assembly of the gray fox (*Urocyon cinereoargenteus*) confirms the basal loss of *PRDM9* in Canidae"

### **Supplementary Information**

Human_NP_064612.2	2 SPEKSQEESPEEDTERTERKPMV

+	+	+

TAVPTDPLFAHPKALGYLHPGQG

grayfox_Scaffold_6__1_c	2 aggcagcctgccagtgtcccgcgGTGCGTG Intron 1

cctccacttcacactgatacgag<0-----[71	: 7328]

atccacttttctatgacactaga

Human_NP_064612.2	25	-KDAFKDISIYFTKEEWAEMGDWEKTRYRNVKRNYNALITI

KDAFK+ISIYF+KEEW +MGDWEK RYRNVKRNY ALIT+

AKDAFKNISIYFSKEEWTQMGDWEKIRYRNVKRNYEALITL

grayfox_Scaffold_6__1_c 7326 TAGgaggtaaatatttaggtacaggtgaactaagaaatggcaac

-0>caactaatctatcaaagcatgagaatgagatagaaacttct

cattcactcacccggagaggacggacatgtgagctaagtca

Human_NP_064612.2	65	LRATRPAFMCHRRQAIKLQ

LRA RPAFMC RR AIK Q

G:G[ggt]	LRAPRPAFMCDRRWAIKPQ

grayfox_Scaffold_6__1_c 7452 GGTAACCG Intron 2	CAGGTcagcccgtatgcatgaacc

<1-----[7453	: 7817]-1> tgccgccttgagggctaca

cactactcgtccggccaca

Human_NP_064612.2	85 VDDTEDSDEEWTPRQQVKPPWMALRVEQRKHQK

VDDTEDSDEEWTPRQQ +	L +	K K

VDDTEDSDEEWTPRQQDEEGQEDLEISS!K-PK

grayfox_Scaffold_6__1_c 7877 gggaggtgggtacaccggggcggtgatt2a ca

taacaacaaagccgaaaaagaaatatcc a ca

atctgtttaagatggatggcgacagttt a cg

Human_NP_064612.2	118	GMPKASFSNESSLKELSRTANL

G+P+ +SN+SSLKELS TA L

GIPRVPISNKSSLKELSETAKL

grayfox_Scaffold_6__1_c 7972 GTATAGG Intron 3	AAGgacagcaaaatatagttgagat

<0-----[7972	: 10552]-0>gtcgtctgaacgtaatcaccat

aacggaatcattggagaaacgg

Human_NP_064612.2	140 LNASGSEQAQKPVSPSGEA

LN S EQ QK VS G+

LNTSSPEQGQKSVSLPGKT	-:N[aat]

grayfox_Scaffold_6__1_c10619 caaaacgcgcatgtccgaaAGTACCTC Intron 4	CAGAT

tacggcaagaactctcgac <1-----[10677 : 11300]-1>

gcgtcaggggaagcttaaa

Human_NP_064612.2	159 STSGQHSRLK-LELRKKETERKMYSLRERKGHAYKEVSEPQDDDYL

T+ H	ELR+K E KMYSL+ERKG AY+EVSEPQDDDYL

PTNLTHVLFSNSELRRKNVEVKMYSLQERKGLAYQEVSEPQDDDYL

grayfox_Scaffold_6__1_c11303 caacacgcttatgcaaaagggaataccgaagcgtcggagccgggtc

ccatcatttcacatggaatatatagtaagagtcaaatgacaaaaat

cccctcactccaacgagttaaggtcaaaagctacagccgcgttccc

Human_NP_064612.2	204	CEMCQNFFIDSCAAHGPPT

CE CQ FFIDSC HGPPT

Y:Y[tat]	CEKCQTFFIDSCTVHGPPT

grayfox_Scaffold_6__1_c11441 TGTAAGTG Intron 5	CAGATtgatcattagatagcgcca

<1-----[11442 : 11637]-1> gaagactttaggctagccc

tggtgccctctttttgcta

Human_NP_064612.2	224 FVKDSAVDKGHPNRSALSLPPGLRIGPSGIPQAGLGVWNEASDLPLGLH

FVKDS VDKG PN SAL+LPPGLRI S IPQAGLGVWN+ASDLPLGLH

FVKDSEVDKGQPNHSALTLPPGLRIRTSSIPQAGLGVWNKASDLPLGLH

grayfox_Scaffold_6__1_c11697 tgagagggagccactgcacccgcaaaataaccggcggtaagtgcctgcc

ttaagataagacaacctctccgtgtgccgtcacgtgtgaaccatctgta

tagctaacgaactcacctgcttgacaaacctgtgtaagcgactgagtac

Human_NP_064612.2	273 FGPYEGRITEDEEAANNGYSWLI----------------

FGPYEG+ITEDEEAAN+GYSWL+

FGPYEGQITEDEEAANSGYSWLLRIIYLFPHPLASHIPP

grayfox_Scaffold_6__1_c11844 tgctggcaaggggggaagtttttaaatctccctgtcacc

tgcaagatcaaaaccaggacgttgttattcactccatcc

tcctgcacaatagacccaccgggaacccccttgcccctt

Human_NP_064612.2	296	-TKGRNCYEYVDGKDKSWANWM

TKGRNCYEYVDGKD SWANWM

ITKGRNCYEYVDGKDNSWANWM

grayfox_Scaffold_6__1_c11961 GTAGACT Intron 6	CAGaaagaattgtgggagattgata

<0-----[11961 : 12325]-0>tcaggagaaatagaaacgcagt

ccagacctgtatagtctgacgg

Human_NP_064612.2	317	YVNCARDDEEQNLVAFQYH

YVNCARDDEEQN VAFQYH

R:R[agg]	YVNCARDDEEQN!VAFQYH

grayfox_Scaffold_6__1_c12392 AGGTAAGGG Intron 7	CAGGtgatgaggggca2ggtctc

<2-----[12394 : 13759]-2> atagcgaaaaaa tctaaa

tgctcgtcaggc gctatc

Human_NP_064612.2	337 RQIFYRTCRVI--RPGCELLVWYGDEYGQELGIKWGSKWKKELMAGR

RQIFYRT R	CELLVWYGDEYGQELGIKWGSKWK EL AG+

RQIFYRTGRT!GHQASCELLVWYGDEYGQELGIKWGSKWKSELAAGK

grayfox_Scaffold_6__1_c13817 acattcagaa4gccgatgccgttgggtgcgcgaatgaataagcggga

gattagcggc gaacggatttgagaaagaatgtagggagagatccga

ggaccactaa ttgcccaggcgcgcgtcgagccggacgggcgcaaga

Human_NP_064612.2	382	EPKPEIHPCPSCCLAFSSQ

EP PEIHPCPSC LAFSSQ

-:A[gca]	EPNPEIHPCPSCSLAFSSQ

grayfox_Scaffold_6__1_c13959 GGTGGGCA Intron 8	CAGCAgcacgacctctttcgttac

<1-----[13960 : 15746]-1> acacatacgccgctctcga

atcagatatacctgccctg

Human_NP_064612.2	401 KFLSQHVERNHSSQNFPGPSARKLLQPENPCPGDQN-QEQQYPDPHSRN

KFLSQH+E NH SQ +P S R+ +P++PCPG QN Q+QQ+ DP	N

KFLSQHLEHNHPSQILPRISVREHFRPKDPCPGCQNQQQQQHSDPQRWN

grayfox_Scaffold_6__1_c15806 atcacctgcacctcacccatgagctccagctcgtcacccccctgcccta

attgaataaaaccattcgtctgaatgcaacgcggaaaaaaaacacagga

acccgtgactcttgccaaataaatcaaatacattgtggggatttagcgt

Human_NP_064612.2	449 DKTKGQEIKERSKLLNKRTWQREISRAFSSPPKGQMGSCRVGKRIMEEE

D+ KGQE KER K L K	QR ISRAFS+P KGQ	C	I++EE

DRAKGQEGKERFKPLPKSIRQRRISRAFSTPCKGQT-TC---EGIVKEE

grayfox_Scaffold_6__1_c15953 gagagcggagatactcaaaacaaatagttactagca at	ggagagg

agcagaagaagtactcagtgaggtcgctcccgagac cg	agttaaa

catatagcaagcatgtataggaatagcttcgcaaaa gt	gaaggag

Human_NP_064612.2	498 SRTG-QKVNPGNTGKLFVGVGISRIAKVKYGECGQGFSVKSDVITHQRT

TG QK+NP +TGKLF GVG++RI +VKY CGQGF+ +S + HQRT

PSTGSQKLNPEDTGKLFKGVGMTRIIRVKYRGCGQGFNDRSHLSRHQRT

grayfox_Scaffold_6__1_c16088 caagtcatacggagattagggaaaaaagatagtgcgtagatccaaccaa

cgcgcaatacaacgattagtgtcgttgtaaggggagtaagcatggaagc

ccacagagtagcacaacggaagaataacatactgaccttgatccatgga

Human_NP_064612.2	546 HTGEKLYVCRECGRGFSWKSHLLIHQRIHTGEKPYVCRECGRGFSWQSV

HTGE YVCRECGRGF+ +++L+IHQR HTGEKPYVCREC GF+ +

HTGENPYVCRECGRGFTHRTNLIIHQRTHTGEKPYVCRECRXGFTERLT

grayfox_Scaffold_6__1_c16235 caggactgtagtgcgtacaaacaaccaacaggactgtagtatgtagata

acgaacatggaggggtcagcatttaagcacgaacatggaggggtcagtc

cgagctttcggtgactacaattcatgaacagggcttcgatgactaggat

Human_NP_064612.2	595 LLTHQRTHTGEKPYVCRECGRGFSRQSVLLTHQRRHTGEKPYVCRECGR

L HQRTHTGEKPYVCR+CGRGF+++S L HQR HTGEKPYVCRECGR

LNEHQRTHTGEKPYVCRKCGRGFTQRS!LSEHQRTHTGEKPYVCRECGR

grayfox_Scaffold_6__1_c16382 cagccaacaggactgtaatgcgtacat1cagccaacaggactgtagtgc

taaaagcacgaacatggaggggtcagc tgaaagcacgaacatggaggg

ccacggacagggcttcaatgactagga ccacggacggggcttcggtga

Human_NP_064612.2	644 GFSRQSVLLTHQRRH

F+++S L THQR H

SFTKRSTLSTHQRTH

grayfox_Scaffold_6__1_c16527 ataaatacaaccaac

gtcagcctgcaagca

ctaggatccacggac

**Figure S1. *PRDM9* alignment output from Genewise**. Alignment using human protein sequence and Urocyon DNA sequence. The four frameshift mutations (indicated by !) a premature stop codon (indicated by X) are highlighted in red. The homology highlighted in yellow was seen across canid species analyzed in previous work (Mooney et al. 2023 & Axelsson et al. 2012).
